# With a little help from my friends: Cooperation can accelerate crossing of adaptive valleys

**DOI:** 10.1101/062323

**Authors:** Uri Obolski, Ohad Lewin-Epstein, Eran Even-Tov, Yoav Ram, Lilach Hadany

## Abstract

Natural selection favors changes that lead to genotypes possessing high fitness. A conflict arises when several mutations are required for adaptation, but each mutation is separately deleterious. The process of a population evolving from a genotype encoding for a local fitness maximum to a higher fitness genotype is termed an adaptive peak shift.

Here we suggest cooperative behavior as a factor that can facilitate adaptive peak shifts. We model cooperation in a *public goods* scenario, wherein each individual contributes resources that are later equally redistributed among all cooperating individuals. We use mathematical modeling and stochastic simulations to study the effect of cooperation on peak shifts in well-mixed populations and structured ones. Our results show that cooperation can accelerate the rate of complex adaptation. Furthermore, we show that cooperation increases the population diversity throughout the peak shift process, thus increasing the robustness of the population to drastic environmental changes.

Our work could help explain adaptive valley crossing in natural populations and suggest that the long term evolution of a species depends on its social behavior.

## 1. Introduction

Adaptive landscapes, introduced by Seawall Wright in the 1930s (Wright 1932), are a useful metaphor for the relationship between genotype and fitness. Under this analogy, fitness is portrayed as a function of genotype, varying between different allele combinations. Complex traits – which depend on two or more loci – can produce rugged adaptive landscapes due to fitness interactions between loci. The simplest instance of a rugged fitness landscape consists of two loci in which mutations are jointly beneficial but separately deleterious. Adaptive valley crossing, also known as an adaptive peak shift, has long been an evolutionary conundrum: how can a population evolve to a higher fitness optimum if it has to “cross” a less fit genotype on the way? Two main stages are needed for such an evolutionary process (Michalakis and Slatkin 1996). First, the fitter genotype must appear in the population. This could happen as a result of sequential mutations in the same lineage, by recombinant offspring of mutant parents or by migration from another population. Second, the fitter genotype will have to spread in the population. However, these two stages have opposing optimal conditions (Weinreich et al. 2005). Mild selection facilitates the first stage of the process: single mutants are more likely to survive in a mild selection regime, increasing the rate of appearance of the fitter genotype by recombination or by acquisition of an additional mutation. In contrast, the second stage is impeded when selection is mild because the fixation probability of a rare genotype decreases (Eshel 1981).

In contrast to Wright’s theory, some researchers have suggested that adaptive landscapes, which can be extended to various topologies in multidimensional genotype spaces (Kingman 1978; Kauffman and Weinberger 1989; Neidhart et al. 2014), might most commonly be single peaked (Bennett 1983; Whitlock 1995, 1997; Gavrilets 2004). However, these theories cannot account for all adaptive landscape topologies, and do not exempt evolutionary biologists from a characterization of evolution on a given rugged adaptive landscape.

Sewall Wright himself was the first to offer a theoretical solution to the adaptive peak shift problem (Wright 1932). His solution, the *Shifting Balance Theory*, was based on a subdivision of the population into small demes in which random genetic drift can increase the frequency of single deleterious mutants. Then, the second mutation can appear on the background of the single mutant. Finally, migration and natural selection can allow the new genotype to expand to other demes. In spite of its novelty and intuitive nature, the *Shifting Balance Theory* has been criticized for the limited range of realistic peak shift scenarios it explains (Coyne et al. 1997, 2000). Additionally, considerable amounts of research focused on finding unique conditions which can facilitate adaptive valley crossing: Dividing the population into smaller subpopulations connected by migration was shown to increase the rate of adaptive valley crossing, even without Wright’s assumptions of increase and decrease in deme sizes as a function of the beneficial genotypes inhabiting them (Bitbol and Schwab 2014); Furthermore, dividing the population into only two populations connected by migration, but changing the selection pressure on each population, was also found to substantially reduce the waiting time for a peak shift (Hadany 2003); Mutation or recombination rates that increase with low fitness were shown to facilitate peak shifts, as this entails that less fit individuals can more rapidly adapt and traverse the fitness valley (Hadany and Beker 2003; Ram and Hadany 2014); Finally, assortative mating was also found to increase the rate of adaptive peak shifts in diploid populations, as it increases the mating between individuals of the advantageous genotype, thus preventing recombination from breaking advantageous allele combinations (Williams and Sarkar 1994). Further theoretical research by Weissman et al. has established the rate of adaptive valley crossing for sexual (Weissman et al. 2010) and asexual (Weissman et al. 2009) populations under different ranges of evolutionary forces such as selection, mutation, recombination and population size. Here we consider an additional factor that can affect the process of crossing adaptive valleys: cooperative behavior. We show that for a population subdivided into demes and connected by migration, cooperation between individuals within the same deme can considerably increase the rate of adaptive peak shifts.

We focus on a *public goods* form of cooperation (Kagel and Roth 1995): all individuals within a deme contribute some resources (thus contributing fitness) to other deme members, and all deme members receive an equal amount of the redistributed resources. This cooperative behavior reduces the fitness difference between different genotypes and therefore effectively "smooths" the landscape to some degree. As a result, less fit mutants are more likely to survive with cooperation, increasing the rate of appearance of multiple mutants. Cooperation has an opposite effect on the fixation of the fittest genotype. Precisely because cooperation smooths the adaptive landscape, it reduces the relative advantage of the fittest genotype, and with it, its fixation probability. Nevertheless, we show that for intermediate levels of cooperation, the increase in the rate of appearance of the fittest genotype outweighs the decrease in its fixation probability, and altogether shortens the total adaptation time. Furthermore, we find that smoothing the adaptive landscape serves the cooperative population in another sense: it increases the population diversity. This increase in diversity is beneficial in evolutionary terms, as it can help populations to overcome environmental changes, parasites etc. (Clarke 1979).

Overall, our results show that cooperation affects adaptive peak shifts substantially and might be an important and overlooked component of complex adaptation.

## Model

We model a population of sexually reproducing haploid individuals, containing two bi-allelic loci. The *ab* genotype is the wild type with fitness 1; the fitness of the single mutant genotypes, *Ab* and *aB*, is 1 − *s*; the fitness of the double mutant *AB* is 1 + *sH*. *s* and *H* are the selection coefficients of the single mutant and the relative advantage of the double mutant, respectively (*s* >0, *H* ≥ 1). We assume equal forward and backward mutation rates for both loci (defined in units of mutations per generation per locus), and denote them by *μ*. Recombination occurs with rate *r* per generation per loci pair.

We model a population composed of *n* demes connected by migration, each of size *k*. Thus the total size of the population is *N* = *n* · *k*. Each generation, individuals of the same deme cooperate. We model cooperation using a *public goods game* (Kagel and Roth 1995): Each individual in the deme contributes a constant fraction *c* of its fitness to the deme (0 ≤ *c* ≤ 1), further referred to as the ‘cooperation level’; the contributions are multiplied by a constant *b* and summed (*b* ≥ 1); finally, this sum is equally redistributed between the deme members. Hence, *c* is the cost of cooperative behavior whereas *b* determines the fold-increase of contributed resources due to cooperation. After cooperating, individuals migrate to other demes with probability *m*. Setting *m* = 1 − 1 / *n* defines a population in which offspring are uniformly distributed among all demes in every generation; setting *m* = 0 determines each deme to be an effectively isolated population. For simplicity, we assume that migration to each deme is equiprobable. Finally, mating occurs between individuals of the newly formed demes. The offspring generation replaces the parent generation, so that population size remains constant and generations do not overlap.

**Table 1.** Parameters used in the analytical model and the stochastic simulation.

| Parameter | Description |
| --- | --- |
| $n$ | Number of demes |
| $k$ | Size of each deme |
| $\mu$ | Mutation rate |
| $r$ | Recombination rate |
| $s$ | Selection coefficient |
| $H$ | Double mutant advantage |
| $c$ | Fraction of resources contributed |
| $b$ | Contributed resources multiplier |
| $m$ | Migration rate |

Fitness is determined by an individual’s genotype and the effect of cooperation. We denote *ω_i_*, as the initial fitness of individual *i*, determined by his genotype (*ab, Ab, aB, AB*), and derive the fitness *ω_i,D_* of individual *i* in deme *D*, after considering cooperation:

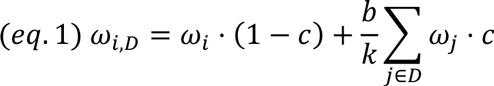

Within this framework, we first analyzed the expected waiting time to a peak shift when each deme contains two individuals (*k* = 2) and the population is fully mixed (*m* = 1 − 1 / *n*) using Branching processes analysis (Eshel 1981) (see Supplementary Information 1). Then, we developed a Wright-Fisher stochastic simulation to account for general deme sizes (*k*) and migration rates (*m*). Simulations include the effects of natural selection, migration, mating, recombination, mutation and drift (see Supplementary Information 2).

The simulations comprise three stages. In the first stage a population inhabited by wild types evolves towards a mutation-selection balance. In the second stage, we simulate the population until a double mutant appears for the first time. This allows us to estimate the expected time for the appearance of a double mutant. In the third stage, the double mutant either goes extinct or fixates in the population (determined by reaching a frequency of 0.99). From this stage we can estimate the probability that a double mutant will fixate (see Supplementary Information 2). Combining the two measures (expected first appearance time and fixation probability) we can estimate the expected waiting time for the appearance of a double mutant that fixates (Hadany 2003). We compared simulation results to the analytical approximation for a fully mixed population (*k* = 2, *m* = 1 − 1/*n*); they were in close agreement (Supplementary Information 1c).

We denote the adaptation time in a population with cooperation level c by τ_c_. The relative difference between the adaptation time of a cooperative population with cooperation level c > 0 (τ_C_) and a non-cooperative population (τ_0_) is denoted by 
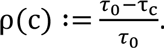

## Results

### A fully mixed population

We start with the special case where demes only have two individuals (*k* = 2) and the population is fully mixed by migration in every generation (*m* = 1 – 1 / *n*). We estimate the relative difference in adaptation time due to cooperation, ρ(c). Fig. 1 presents ρ(c = 0.6), derived from our approximation (see Model and Supplementary Information 1), as a function of the selection coefficient, *s,* and the double mutant advantage, *H*. Higher recombination rates restrict the possibility of a peak shift, as recombination tends to break beneficial gene combinations (Eshel and Feldman 1970). This is consistent for cooperative behavior as well (Fig. 1, compare A and B). Cooperation has contrasting effects over the two main stages of the peak shift. First, it reduces the disadvantage of the single mutants (*Ab, aB*), therefore increasing their survival. As a result, the waiting time for the first double mutant (*AB*) shortens in comparison to a non-cooperative population (Fig. 1). On the other hand, cooperation reduces the benefit of the double mutant, thus decreasing its survival probability. When selection against the single mutant (*s*) or the advantage of the double mutant (*H*) are high (upper right corners in Fig. 1), the decrease in waiting time for the double mutant’s first appearance is more pronounced than the decrease in its fixation probability, resulting in faster adaptation and thus a higher ρ value.

Note that some conditions allow for a peak shift only in a non-cooperative population, whereas a cooperative population is not expected to cross the adaptive valley (Fig. 1, black areas). The public goods cooperation, as modeled here, cannot expand the range of conditions allowing for a peak shift because cooperative behavior effectively decreases the double mutant advantage (see a formal proof for a fully mixed population in Supplementary Information 1b).

**Figure 1.**
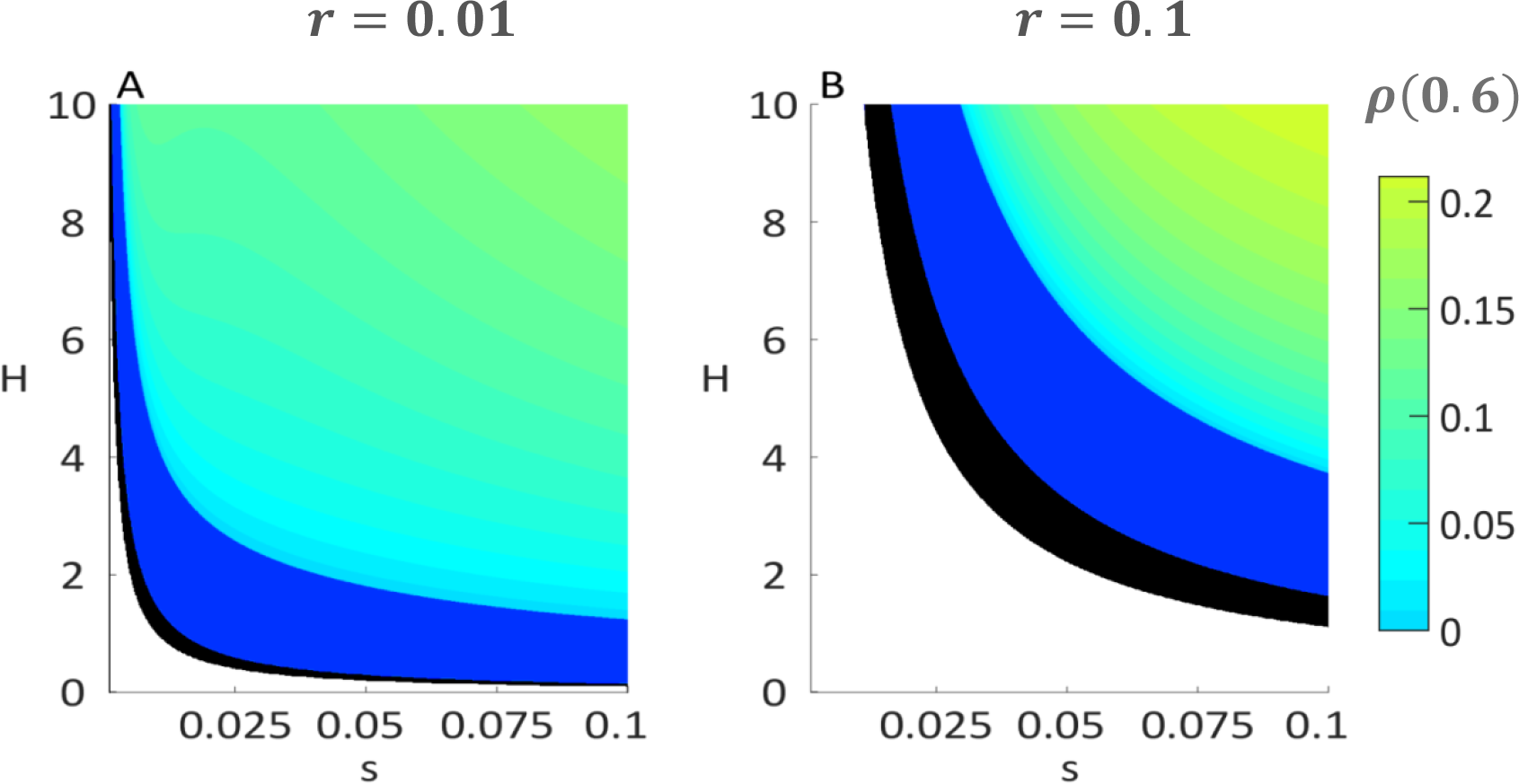
Cooperation affects adaptation time in a mixed population. Results are based on analytical approximations (Supplementary Information 1a). White areas represent parameters where both cooperatives and non-cooperatives are not expected to achieve a peak shift; black areas are parameters for which only non-cooperatives are expected to achieve a peak shift; blue areas represent parameters for which both cooperatives and non-cooperatives achieve a peak shift, but non-cooperatives accomplish the process faster. Teal to green areas are parameters for which cooperatives achieve peak shift faster than non-cooperatives, and the colors represent the relative difference in the expected time to peak shift due to cooperation (ρ). Panels A and B present the results for recombination rates *r* = **0.01,0.1**, respectively. Additional parameters are: N = 10,000, μ = 10^−5^, c = 0.6, b = 1.2.

### A subdivided population

Next we analyzed the rate of adaptation in populations divided into demes containing more than two individuals (*k* > 2). The division to larger demes changes the frequencies of single and double mutants, rendering our approximation no longer compatible for the multi-level selection between and within demes. Thus, all further analysis is based on stochastic simulations (see Model).

First, we examine the effects of selection (*s* and *H*) on the adaptation time with and without cooperation. Fig. 2 shows the relative difference in adaptation time due to cooperation, ρ, for populations divided to demes that contain 10 individuals. Similarly to Fig. 1, *ρ* is a function of the selection coefficient, *s*, on the horizontal axis, and the double mutant advantage, *H*, on the vertical axis. The division of the population to demes retains the properties exemplified for a mixed population; high recombination values narrow the range of conditions leading to a peak shift (Fig. 2 compare A and B), and cooperation cannot extend the parameter range enabling the population to cross the adaptive valley (Fig. 2B, black area).

However, cooperation in a subdivided population accelerates complex adaptation even more than in a fully-mixed population; i.e., the subdivided population attains higher p values (note the red-yellow hues of Fig. 2 in comparison to Fig. 1; both figures have the same color scales).

**Figure 2.**
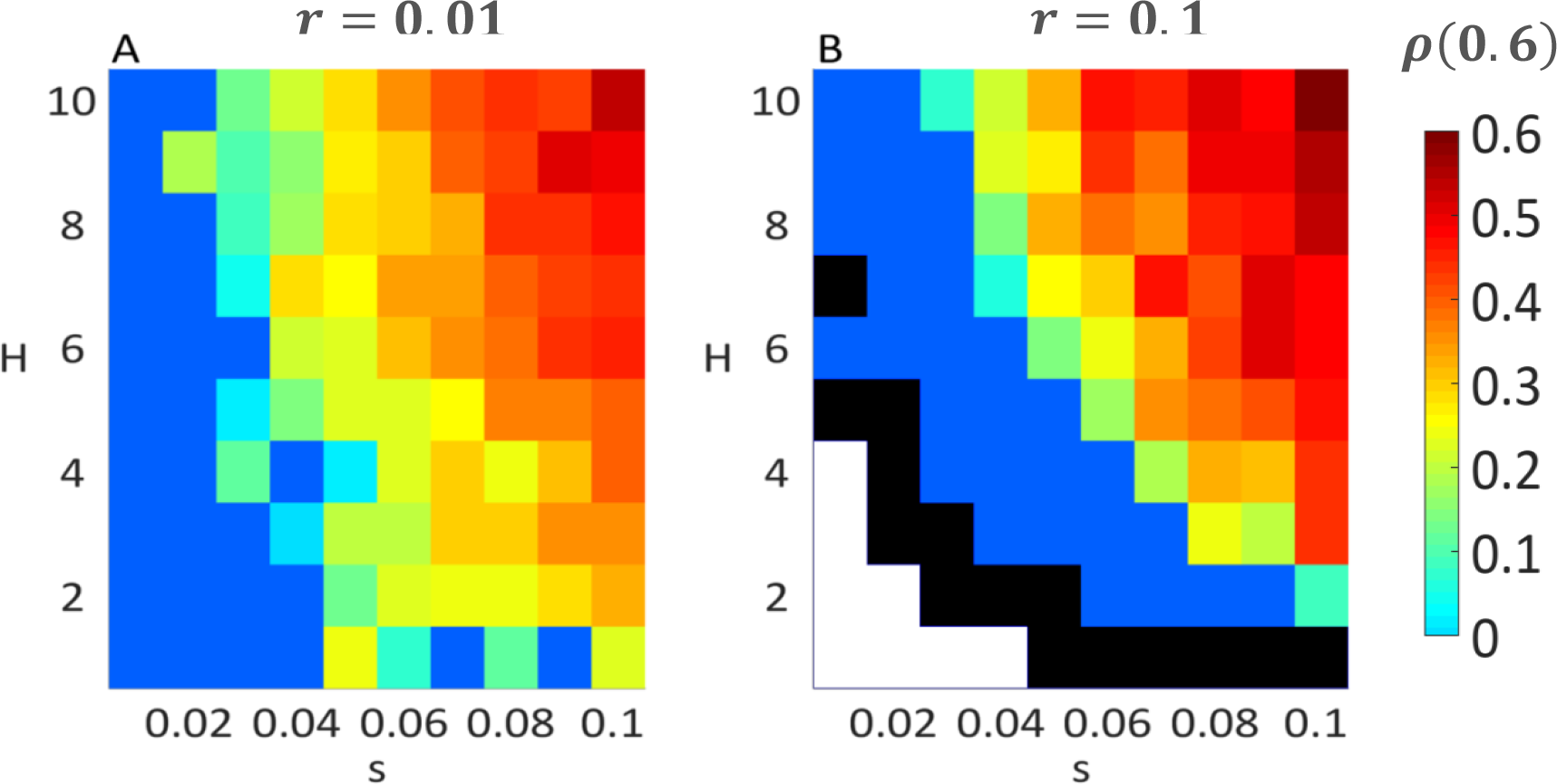
Cooperation affects adaptation time in a subdivided population. Results are based on stochastic simulations (Supplementary Information 2). The x-axis represents the selection coefficient, *s*, and the y-axis represents the double mutant advantage, *H*. White areas represent parameters where both cooperatives and non-cooperatives are not expected to achieve a peak shift; black areas are parameters for which only non-cooperatives are expected to shift a peak; blue areas represent parameters for which both cooperatives and non-cooperatives achieve a peak shift, but non-cooperatives do so faster. Teal to red areas are parameters for which cooperatives achieve a peak shift faster than non-cooperatives, and the color represents the average relative difference in the expected time for a peak shift due to cooperation, ρ, averaged over ≥ 1200 simulations per parameter set. Panels A and B represent results for low and high recombination rates: *r* = 0.01,0.1, respectively. Additional parameter values are *n* = 1,000, *k* = 10, μ = 10^-5^, *c* = 0.6, *b* = 1.2, *m* = 0.01.

We investigated the effects of the cooperation levels, *c*, and the cooperation benefit, *b*, on the fixation of double mutants. In Fig. 3 we break down the dynamics of the double mutant fixation to the waiting for the appearance of a double mutant (Fig. 3A), the fixation probability of a double mutant (Fig. 3B), and the overall time to adaptation (Fig. 3C). For higher benefit produced from contributed resources, *b* values, the curves of first appearance and fixation probability become steeper (Fig. 3A and B). Higher *b* values effectively diminish the influence of each genotype’s fitness and increase the influence of the pooled fitness. Thus, increasing *b* "smooths" the adaptive landscape and decreases the waiting time for the appearance of a double mutant (Fig. 3A). On the other hand, increasing *b* reduces the advantage of the double mutants and therefore decreases its fixation probability (Fig. 3B). Although high cooperation levels, *c* values, substantially shorten the waiting time for appearance of a double mutant, they also reduce its fixation probability. Therefore, intermediate cooperation levels minimize the adaptation time by striking a balance between shortening the waiting time for appearance of a double mutant and decreasing its fixation probability (Fig. 3C).

Of course, when the population is non-cooperative (*c* = 0), *b* doesn’t affect the results. With full cooperation (*c* = 1), all individuals within a deme have the same fitness regardless of their genotype, thus yielding the same relative fitness for all *b* values (eq. 1).

**Figure 3.**
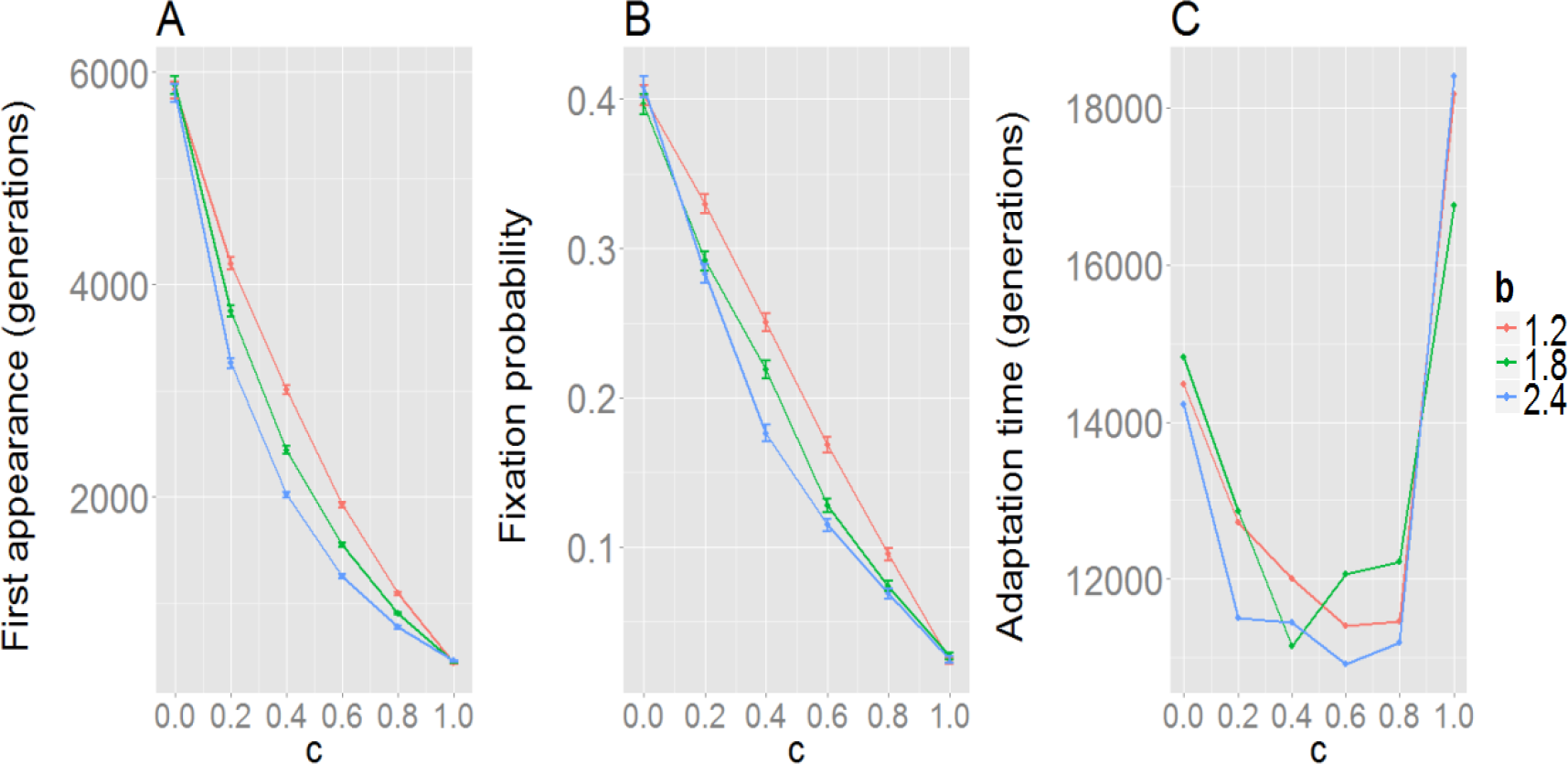
Effect of cooperation on complex adaptation in a subdivided population. The figure shows simulation results for **(A)** first appearance of a double mutant; **(B)** fixation probability of a double mutant after appearance; and **(C)** total adaptation time. Markers show averages of ≥ 5,000 simulations; bars show standard error of the mean. Colors indicate different *b* values (red: 1.2, green: 1.8, blue: 2.4). Additional parameter values: r = 0. 01, n = 1,000, k = 10, μ = 10^−5^, m = 0.01, s = 0.05, H = 5.

### Population diversity

Another interesting facet of the evolutionary process of peak shifts is the population diversity. A genetically diverse population is more robust to environmental changes that change genotypes’ fitness, and is thus less likely to go extinct due to such changes (Clarke 1979). In order to measure the genetic diversity we use Shannon’s Index, normalized by the number of genotypes:

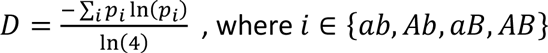

where *p_i_* is the frequency of genotype *i* in the entire population. *D* ranges from zero to one, indicating only one genotype exists or that all genotypes are found in the population in equal frequencies, respectively.

We define the relative increase in diversity between cooperative and noncooperative populations to be 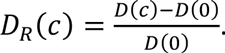. For diversity analysis, we simulated the peak shift dynamics for 100,000 generations and recorded the diversity every 100 generations. In Fig. 4 we show the difference in diversity between cooperative and non-cooperative populations, *D_R_*(*c*), against varying *s* and *H* values. For all examined selection coefficients, *s* and *H*, cooperation increases the diversity relative to a non-cooperative population, usually by more than two-fold (*D_R_*(0.6) > 1, Fig. 4A).

During a peak shift, populations reach mutation-selection balance under two selective regimes: First before the successful double mutant appears and second after it fixates. The population diversity in the second mutation selection balance is lower than in the first, since the selective disadvantage of the single mutant compared to the double mutant is higher than compared to the wild type. Cooperation reduces the effective disadvantage of single mutants as well as the advantage of double mutants, and therefore increases diversity both before and after the peak shift (Fig. 4; Supplementary Information 3). However, when selection is very weak, this effect diminishes (see Fig. 4, bottom-left corner). When selection is strong, cooperators achieve a peak shift faster and spend more time in the second mutation-selection balance, relative to non-cooperators (Supplementary Information 3). Hence, strong selection diminishes the diversity advantage of cooperators (Fig. 4A). Overall, we see that a cooperative population retains, on average, higher diversity than a non-cooperative population during the entire peak shift process (Fig. 4B, Supplementary Information 3).

**Figure 4.**
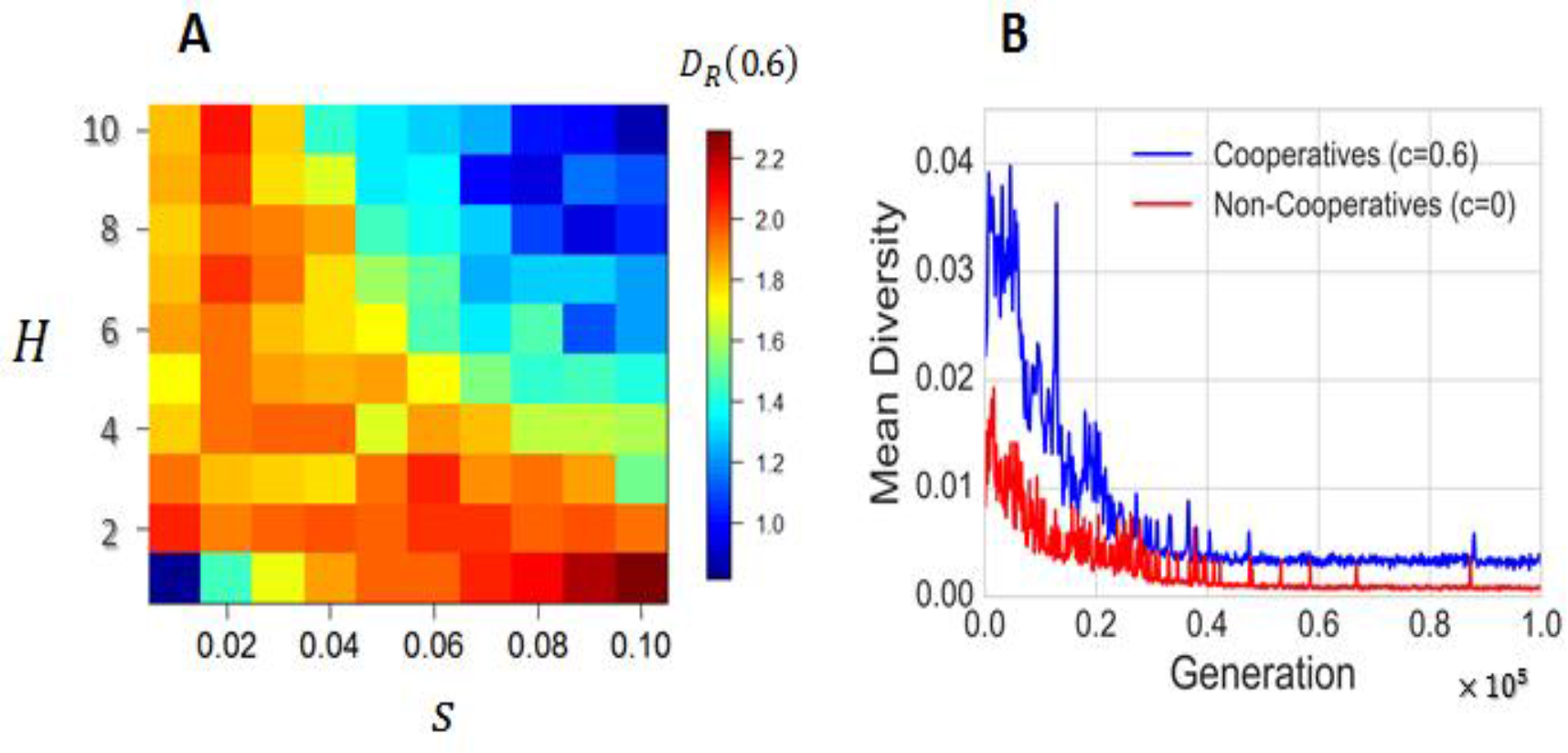
Effect of cooperation on genetic diversity during a peak shift. **(A)** The relative increase in diversity, *D_R_*(0.6), as a function of the selection coefficient, *s*, and the double mutant advantage, *H*. Cooperating populations are more diverse throughout the parameter range. Data is averaged over ≥ 200 simulations per parameter set. **(B)** The average genetic diversity (taken over ≥ 200 simulations) in a cooperative population (blue line) is higher than that of a non-cooperative population (red line) for selection coefficient *s* = 0.05 and double mutant relative advantage *H* = 5 at almost every time point. Additional parameter values are: n = 1,000, k = 10, μ = 10^−5^, r = 0.01, c = 0.6, b = 1.2, m = 0.01.

Although there is a marked increase in diversity for a cooperative population, the dynamics of the double mutant’s fixation are similar in cooperative and noncooperative populations. A peak shift can occur either by a rapid takeover of double mutants occupying entire demes, or by a gradual increase of the number of double mutants in each deme. We find that the maximal variance of double mutants among demes during the peak shift is similar with or without cooperation. However, the duration of the double mutant takeover in a cooperative population appears to be more variable than in a non-cooperative one (See Supplementary Information 4), suggesting that a metapopulation composed of cooperative populations would be highly variable over a long period of time.

### Discussion

In this study we have shown that cooperation can be an important factor in the evolution of complex genotypes. We have found that in a *public goods* scenario, cooperative behavior can accelerate a peak shift, relative to non-cooperative behavior. However, in our model, cooperators can only achieve a peak shift under conditions enabling so for non-cooperators. Our results indicate that the adaptive advantage of cooperative behavior increases with the strength of selection, and that the range of conditions where cooperation is beneficial is expanded for low recombination rates. The effect of cooperation on fixation time is usually not monotonous, and intermediate values of cooperation (i.e. not a full investment of one’s resources in cooperative behavior) might be optimal for achieving peak shifts. Additionally, we examined how cooperation affects genetic diversity. Cooperation smooths the adaptive landscape by decreasing selection intensity, and therefore increasing genetic diversity before, during and after a peak shift (Fig. 4 and Supplementary information 3).

Cooperation is a widespread biological phenomenon, with a vast body of theoretical literature supporting its evolutionary feasibility (Hamilton 1964; Trivers 1971; Axelrod and Hamilton 1981; Nowak 2006; West et al. 2007b). The focus of our work is the effect of cooperation on the dynamics of complex adaptation, rather than the conditions leading to the emergence of cooperative behavior. Importantly, we note that our results hold for population with low relatedness (in a fully mixed population with migration rate *m* ≈ 1) as well as populations with high relatedness (*m* ≪ 1). Our model assumes that migration is equal between all demes (equivalent to spatial homogeneity between the demes). This assumption can be violated if migration is fitness-associated at the individual level. In this case, less fit individuals may be inclined to migrate more often in order to improve their offspring’s genotypes (Gueijman et al. 2013) and cooperative populations would have lower effective migration rates than non-cooperative ones. However, we do not expect this to have a qualitative effect, because the advantage of cooperative populations does not depend on the level of relatedness.

Recently, Komarova has shown that peak shifts in asexual populations in a spatially heterogeneous environment can be facilitated by cooperation, when genes affecting cooperation also determine the fitness, and cooperators directly compete with non-cooperators (Komarova 2014). However, we model the loci determining the fitness as independent of cooperation, and the results are relevant for peak shifts of genes that are not directly affected by cooperation. Furthermore, we show that this can occur for sexual organisms, and even without explicit spatial constraints (*e.g.* fully mixed populations; Fig. 1).

Interestingly, cooperative behavior in our model does not change the maximal between-deme variance of the double mutant distribution (Supplementary information 4). This implies that the advantage of cooperation does not stem solely from increased genetic drift. Rather than increasing genetic drift, or bypassing the adaptive peak, like Wright and Fisher have suggested (Wright 1932; Bennett 1983), cooperation directly changes the intensity of selection. Nevertheless, cooperation changes the variance of the double mutants’ duration of fixation (Supplementary information 4). If different runs of our simulations are viewed as possible outcomes of independent populations under the same conditions, then the time needed for a double mutant to spread in these cooperative populations varies substantially. That some populations lag behind in crossing adaptive valleys might result in competition between populations or even lead to speciation. The added advantage due to high genetic diversity in cooperative populations can also influence their survival probability. Populations might encounter new parasites, predators, or abiotic environmental changes, against which some of the intermediate genotypes might have an advantage (Benton 2009). Maintaining intermediate genotypes could in such cases be a substantial advantage of cooperation.

Multicellular, sexually reproducing organisms are an obvious fit to our assumptions, if recombination rates between the relevant loci are low and selection is not too weak. Our model can also be relevant to bacteria, for example, which can face a peak shift challenge when developing antibiotic resistance (Salverda et al. 2011; de Visser and Krug 2014). Furthermore, some mutations that confer antibiotic resistance carry a fitness cost, but can be compensated by additional mutations that are beneficial in the presence of antibiotics and slightly deleterious in its absence (Andersson and Hughes 2010). Bacteria carrying both resistance and compensation mutations in an environment currently without antibiotics would need to cross an adaptive valley to become non-resistant and uncompensated. Although bacteria do not reproduce sexually, they can perform some horizontal gene transfer (Ochman et al. 2000; Thomas and Nielsen 2005), as befitting our model. Bacterial cooperation is also documented: bacteria often aggregate to produce biofilms or molecules that can be considered as *public goods* (Kreft 2004; West et al. 2007a; Nadell et al. 2008). Our results suggest that cooperative bacteria may enjoy an additional benefit of crossing adaptive valleys faster and having increased genetic diversity. Such knowledge on bacterial population dynamics might be used to devise strategies to fight antibiotic resistance, as other evolutionary processes are used for predictions of efficient treatment strategies (Obolski and Hadany 2012; Obolski et al. 2015; Perron et al. 2015; Caudill and Wares 2016; Obolski et al. 2016).

To conclude, we suggest a possible interplay between evolutionary forces and social behavior: Cooperative behavior can hasten the appearance of complex genotypes with increased fitness, which in turn might play a role in maintaining cooperative populations.

## Acknowledgements

This project was supported by the Israeli Science Foundation 1568/13 (LH), the Minerva Center for Lab Evolution (LH) and by a fellowship from the Manna Program in Food Safety and Security (UO).

## Supplementary Information

### Supplementary Information 1

#### <u>1a.</u> <u>Analytic approximation</u>

Using Branching processes (Harris 1948) we analyze the probability of a peak shift for the case of *k* = 2 and 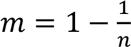. We define *ω_x_y_* to be the fitness of genotype x, when genotype y is its partner.

Since the fitness of *Ab* and *aB* is the same, and so is their contribution to their partners we will denote *sm* (single mutant) as either *Ab* or *aB*.

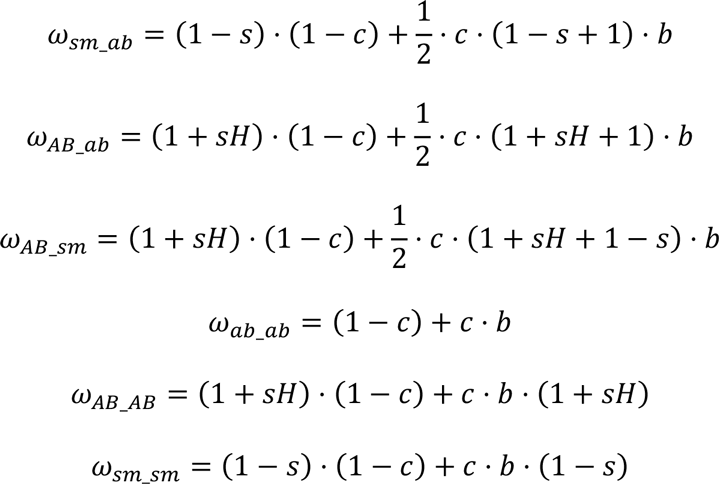

Since *ab* is the wild type and is the most common genotype, and since the population is fully mixed, most cooperating couples are of two *ab* individuals. Therefore the fitness of most individuals is *ω_ab_ab_*.

We normalize the fitness of all individuals relative to the most common fitness ω*_ab_ab_* and define:

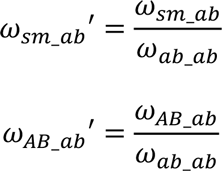

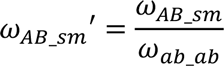

By neglecting terms of the order of *μ*^2^ we can say that the partner of an *sm* individual, at the mutation-selection balance, would be *ab*. Now we can calculate *s*’, the effective selection coefficient when taking cooperation into account:

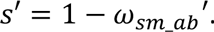

We can approximate the probability *q* that an *AB* individual would be formed in the next generation, given no other *AB* individuals exist in the current generation:

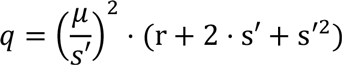

Hence, the probability that the first *AB* individual would appear in the population at a certain generation, given that it had not appeared earlier is:

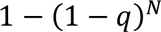

which can be approximated by *q* · *N*, where *N* = *n* · *k* is the size of the population. The expected time for appearance of the first *AB* individual (*T_first_*), which is geometrically distributed, is: 
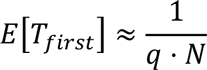

The approximation of the average fitness, marked by 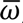, is:

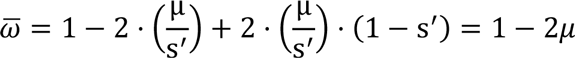

The progeny of an *AB* individual, marked by *p*, is:

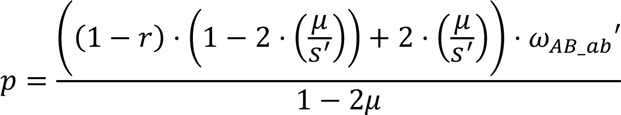

Therefore, if we assume that the number of offspring is Poisson distributed and its mean is slightly above one (Eshel 1981; Hadany 2003), the probability that an *AB* genotype would fixate in the population, rather than go extinct, denoted by π, is approximated by:

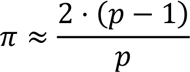

Thus we can approximate the expected time for appearance of an *AB* individual that will go to fixation by:

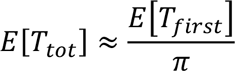

#### <u>1b.</u> <u>Expected progeny analysis for partners:</u>

If the fitness of an *AB* individual is 1 + *sH*, and μ ≪ *s*, then the expected number of the *AB*’s offspring which are themselves *AB* is approximated by:

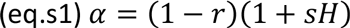

The fitness of an *AB* when its partner is *ab*:

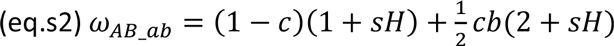

The fitness of a *ab* when its partner is *ab*:

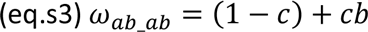

We normalize the new fitness

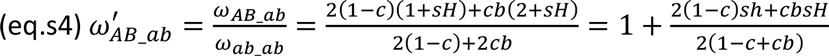

Using (eq.s4) and (eq.s1) we get:

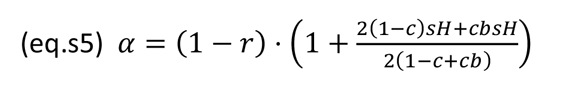

In order for *AB* to fixate, we require that *α* > 1.

Since 0 ≤ *c* ≤ 1 and *b* ≥ 1 we get that the denominator in (eq.s5) 2(1 − *c* + *cb*) is always positive and therefore we can multiply (eq.s5) and get:

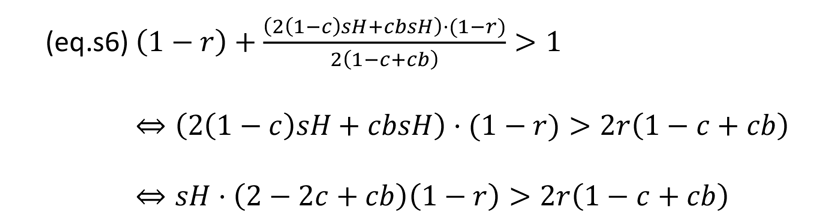

Since *r* < 1 and *c* ≤ 1, we have that (2 − 2*c* + *cb*)(1 − *r*) > 0 and therefore

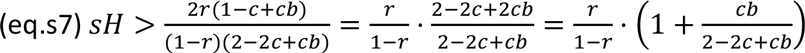

Differentiating the right hand side of (eq.s7) with respect to *c* yields:

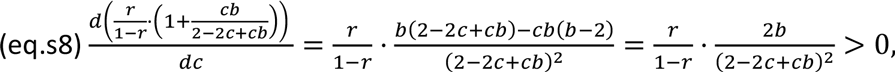

Therefore 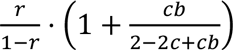 is monotonically increasing in *c* for all *b* ≥ 1, and 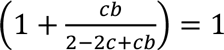 only when *c* = 0.

**Thus, any increase in *c* restricts the parameter range enabling *AB*’s fixation.**

#### <u>1c.</u> <u>Comparison between analytical approximation and simulation results</u>

In order to verify our approximations we used stochastic simulations. Fig. S1 shows the results of the simulation (blue line) and the approximation (red line), for various cooperation levels. This is shown for the time of first appearance of the double mutant (Fig. 2A) and the double mutant’s fixation probability (Fig. 2B), as a function of the cooperation level (*c*). When *c* = 0, we are reduced to the results presented, and verified against simulations, in (Hadany 2003). We can see that our approximation is accurate when *c* increases, but it does remain slightly biased.

Note that both the waiting time for appearance of the double mutant and its fixation probability decrease with *c*.

**Figure S1.**
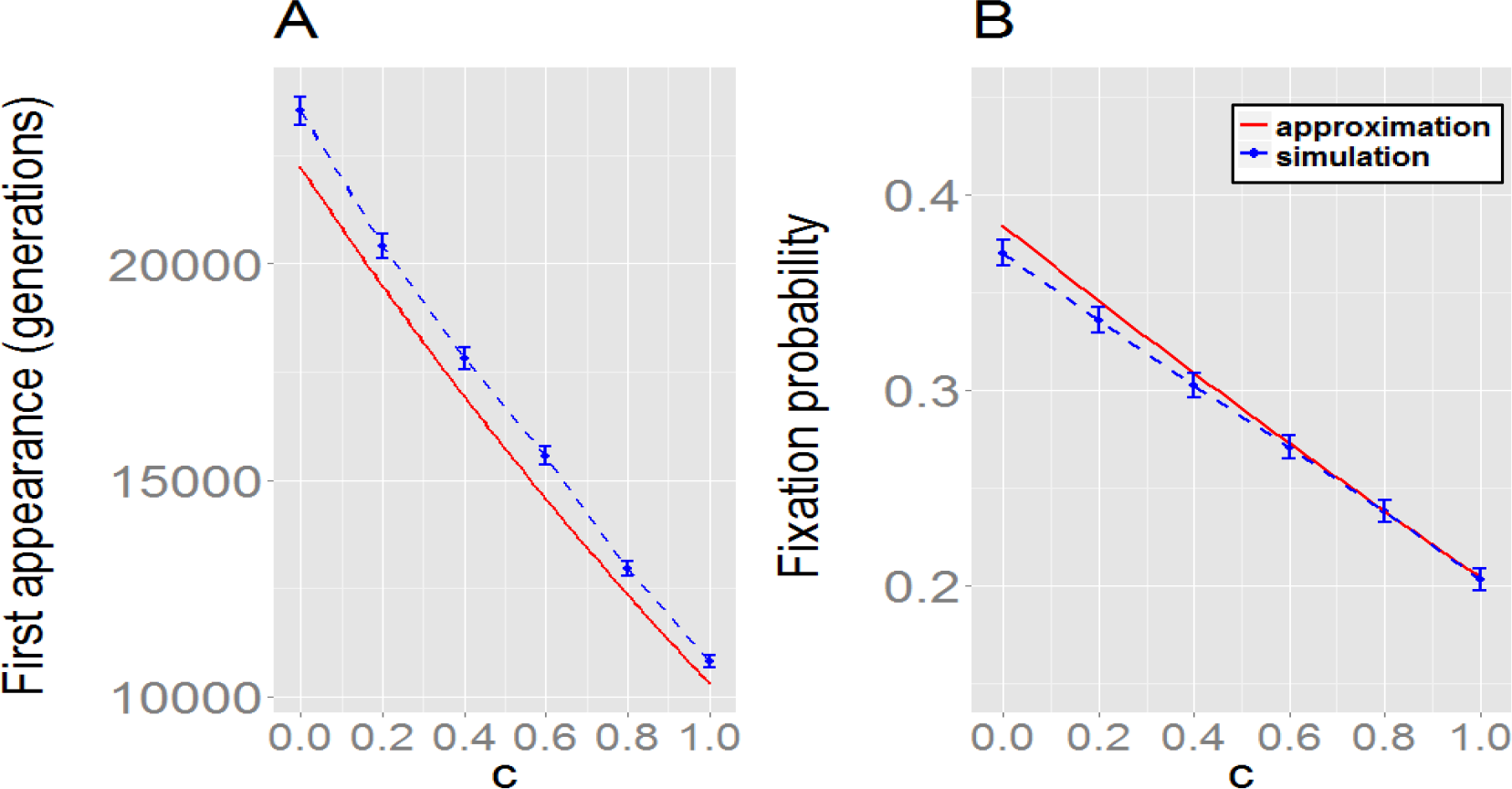
Comparison between simulation and analytical approximation in a fully mixed population. We plot the effect of cooperation on the first appearance of a double mutant and its fixation. As the level of cooperation increases (x-axis) both the waiting time for appearance of a double mutant (A) and the fixation probability of a double mutant (B) decrease. We can see that the analytical approximation and the simulation results are in close agreement. Other parameters are: *s* = 0.05, *H* = 5,*n* = 5,000, *k* = 2, 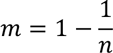 μ = 10^−5^, *r* = 0.01, *b* = 1.2.

### Supplementary Information 2 – Simulation design

The simulation is composed of a genotype frequencies vector, representing the structure and composition of the population, and of several functions, each representing a different process: selection, migration, mating and recombination, mutation and drift. One generation ends after all functions have been applied and the population is replaced.

We initialize the simulation with a population comprised solely of wild type individuals. Every generation, we apply the selection function and calculate the fitness separately for each deme. We calculate the ‘donation pool’ of every deme and then derive the new fitness of each individual by (eq. 1) in the main text:

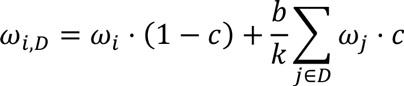

After that we apply the migration function to the fitness-weighted frequencies. The frequency of genotype *i* in deme *D*, after selection and migration (marked as 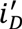) is:

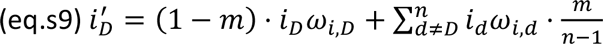

After selection and migration, we apply the mating-recombination function. We calculate the frequencies of the genotypes in the next generation, within each deme, according to the frequencies in the parent generation, and the recombination rate, *r*. Finally, we apply mutation.

Note that we do not normalize the results after applying the selection function. Thus, demes with higher mean fitness would export more individuals.

The last phase in each generation is drift, simulated by drawing *k* random numbers from a multinomial distribution based on the relative genotype frequency in each deme, and repopulating the demes. We then transform the quantities of every genotype back into frequencies. After applying all the functions, and initiating the new frequency vector, the simulation proceeds to the next generation.

We start each simulation with running 3,000 generations, beginning with a population composed solely of wild type individuals. At this stage we define the fitness of genotype AB to be 0 in order to reach mutation selection balance while no double mutant is yet formed.

The parameters we use are *n, k, r, μ,s, h, b, c, m*, as defined in Table 1 in the main text. The frequency vector has 4 · *n* coordinates, where 4 coordinates describe the frequencies of the different genotypes in a specific deme. For instance, for the population vector *V*, the frequencies of genotypes *ab,Ab,aB,AB* in deme *j* are [*V*(4*j*),*V*(4*j* + 1),*V*(4*j* + 2),*V*(4*j* + 3)].

#### 1. Fitness

For deme *k* we calculate the pool of fitness donations using matrix product:

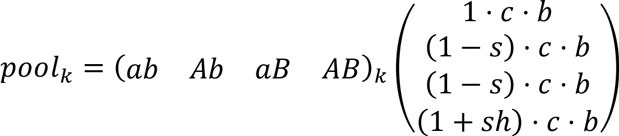

After calculating the pool we can calculate the new fitness of every individual, and by that, the frequency of every genotype after applying selection. The new frequencies of genotypes in deme *k* would be:

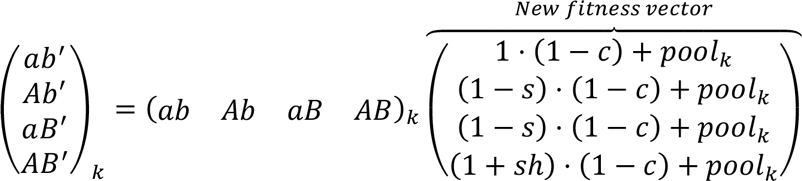

Where the vector (*ab Ab aB AB*)*_k_* represents the frequencies of the genotypes in generation *N*, and the vector (*ab′ Ab′ aB′ AB′*)*_k_* represents the frequencies of the genotypes after selection.

#### 2. Migration

According to eq.s9, we can define a migration matrix that would simulate the change in the population composition due to migration. For instance, if there are 3 demes in the population, the matrix would be defined as follows:

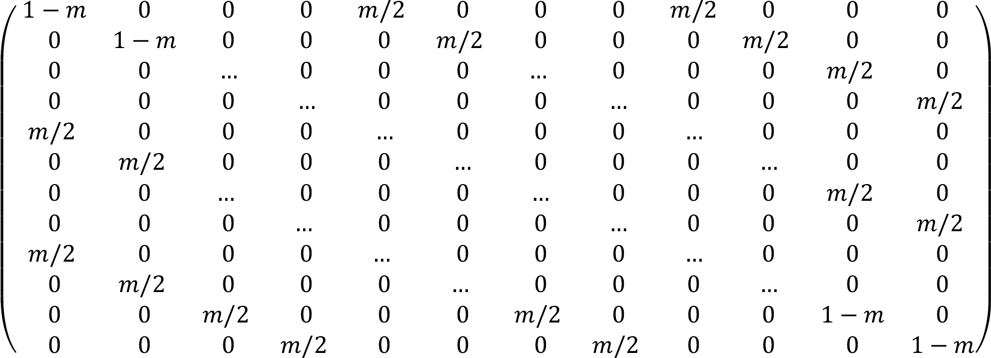

#### 3. Mating and recombination

Let *i* ∈ {*ab,Ab,aB,AB*} be the frequency of genotype *i* in a specific deme in generation *N*, and *i*′ the frequency of genotype *i* in that same deme in generation *N* + 1. By this we get:

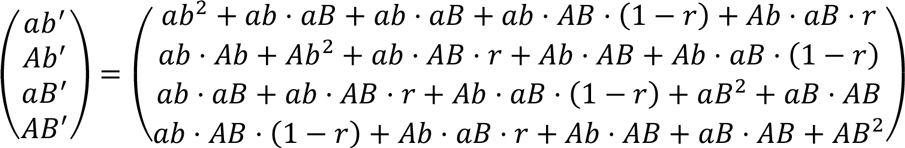

#### 4. Mutation function

The frequency of each genotype after mutation takes place can be written as:

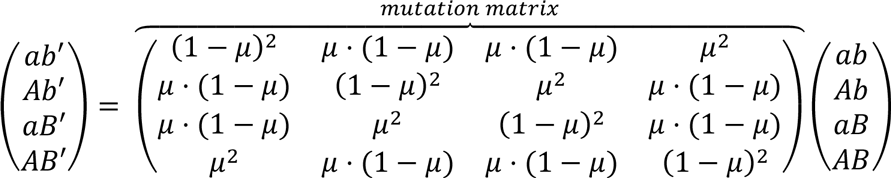

In this notation, *i* ∈ {*ab,Ab,aB,AB*} represent the frequency of genotype *i* in a specific deme before mutation and *i*′ represents the frequency of genotype *i* in that same deme after mutation takes place. Since the mutation occurrences are independent variables, each deme is multiplied by the same matrix. Moreover, the probability of mutation occurrence is homogenous in time; therefore the same matrix is valid for the entire simulation.

Simulations were performed using Python 3.3. When multiple parameter sets were examined, assignments with random selection of the desired parameter sets were sent to a computer cluster until the pre-determined number of runs was satisfied for each parameter set. Hence the statements of ‘at least ## simulations’ found in the main text.

### Supplementary Information 3 – Diversity analysis

In order to see the change in genetic diversity as a function of generations, we plotted the average graph for several parameter sets. For each parameter set, we chose simulations that ended with fixation of double mutants (limited by 100,000 generations) and calculated the average of two time points:

- *T*_1_ – The generation of the first appearance of a double mutant that will eventually fixate (in contrast with appearance of double mutants that appear and then extinct).
- *T*_2_ – The generation in which the double mutants reached a frequency of 0.99 for the first time.

These two time points divide the process to three phases:

1. MSB1 – First mutation-selection balance. The time until the first appearance of a successful double mutant.
2. Takeover – The phase between the first appearance of a successful double mutant and its fixation.
3. MSB2 – Second mutation-selection balance, after the fixation of the double mutants.

For simulation *j* let 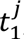, 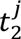 be the time of the first appearance of a successful double mutant and the time of its fixation, respectively. We normalize each generation *i* in simulation *j* to its relative location within its phase in the following manner:

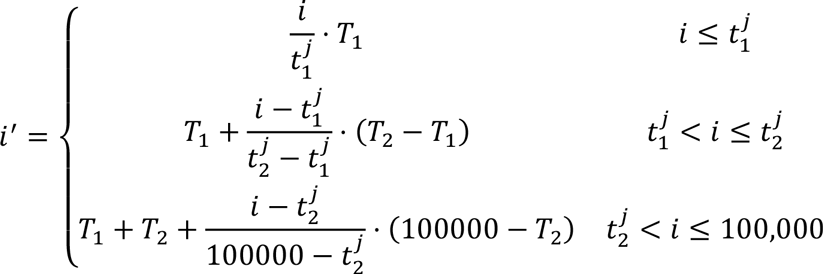

After normalization, we smoothed the diversity of the simulations with a 10 generation moving average and plot the results. In the plots presented below, we show that cooperatives attain higher diversity during the first and the second mutation-selection balances. In addition, during the takeover cooperatives reach higher levels of diversity. We can see that occasionally the diversity in a noncooperative population is higher than in a cooperative population. This happens only for relatively short periods and mostly when the two populations are not well synchronized: For example if the cooperative population has already crossed the valley and is in the second mutation-selection balance (after fixation) while the non-cooperative population is still in the first stage (see s=0.09, H=9 plot; generations ~10,000-20,000). Nevertheless, as shown in the Fig. 4 in the main text, the average diversity in the entire process is almost always higher for cooperative populations.

**Figure S2.**
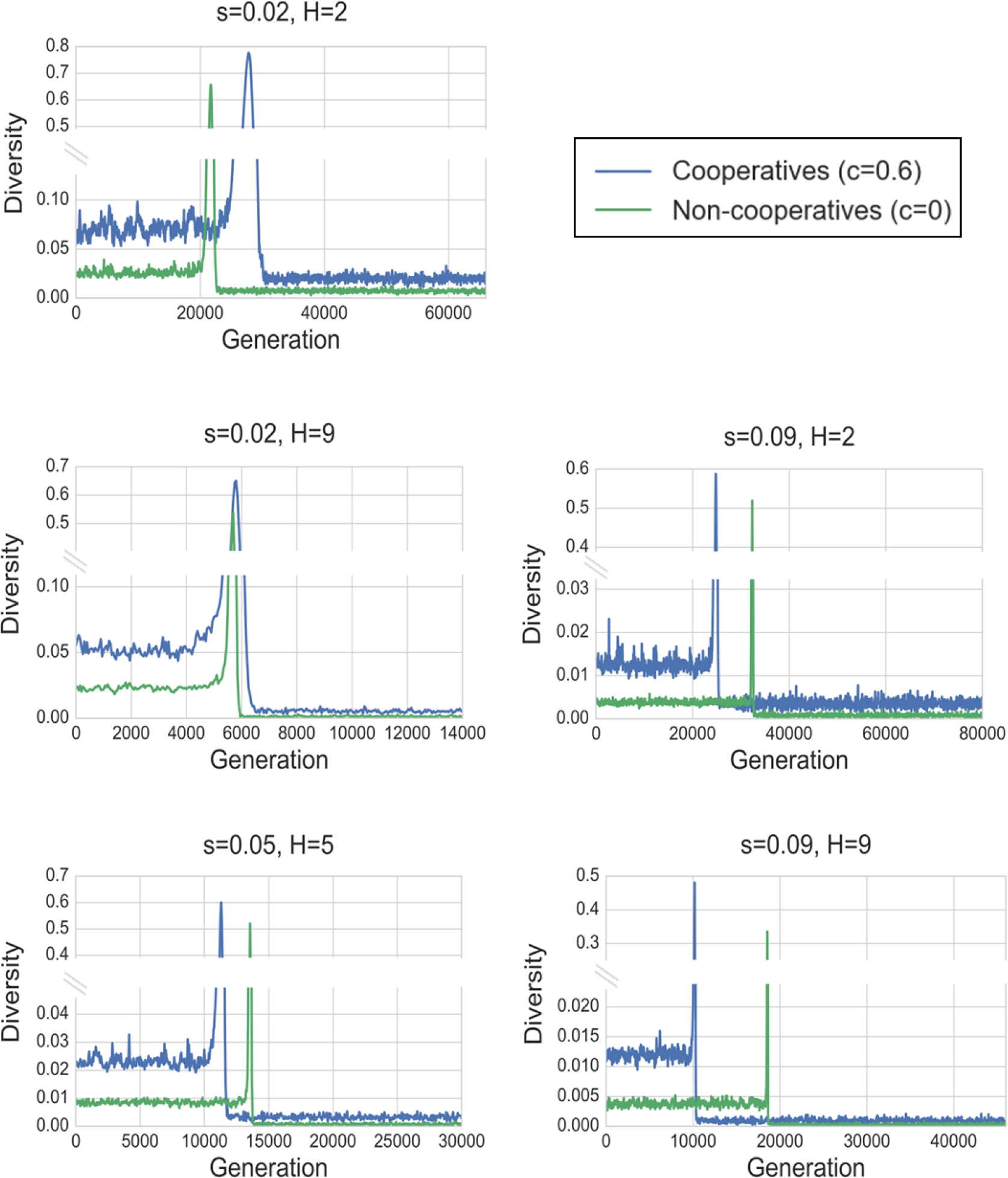
Mean diversity is higher for cooperative population. For each phase (MSB1, Takeover, MSB2) the diversity in a cooperative population is higher than in a non-cooperative population. Each plot is based on at least 180 simulations. Note that the y-axes are broken. Parameters are: *n* = 1,000, *k* = 10, *μ* = 10^−5^, *r* = 0.01, *c* = 0.6, *b* = 1.2

### Supplementary Information 4 – SD of double mutants between demes

In this section we examined the variation in the number of double mutants between demes during the fixation process. A peak shift can occur either by a rapid takeover of double mutants occupying entire demes, or by a gradual increase of the number of double mutants in each deme. The spread of double mutants in the population can be expressed by the variance of the number of double mutants between the demes:

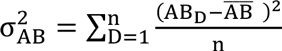

Where AB_D_ is the number of double mutants in deme D, and 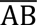 is the average number of double mutants per deme 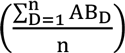.

A population composed of demes with a similar number of double mutants would have low variance, whereas high variance is expected when the population is composed of demes fully inhabited by double mutants and demes completely devoid of them. We find that the cooperation level (c) does not influence the maximum variance attained during a peak shift (compare red to blue curves in the figure), indicating that the peak of the distribution of double mutants among demes during the fixation process is similar with or without cooperation. However, the dynamics of the double mutant spread in a cooperative population are more variable than in a non-cooperative one.

**Figure S3.**
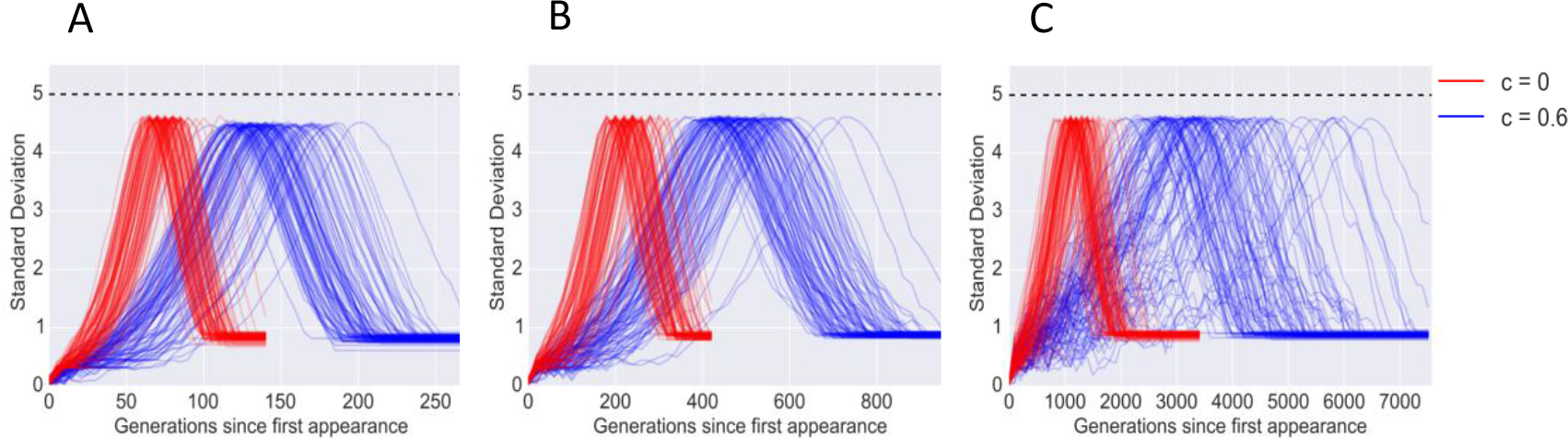
The change in variance of double mutants across demes during a peak shift. Standard deviation of the number of double mutants between demes (*σ_DM_*) is plotted for 100 simulation runs of a cooperative population (*c* = 0.6, blue) and a non-cooperative population (*c* = 0, red), under several sets of selection coefficients: *s* = 0.1, *h* = 10 **(A)**, *s* = 0.05, *h* = 5 **(B)** and *s* = 0.02, *h* = 2 **(C)**. Each simulation begins from the first successful double mutant (a double mutant that fixates in the population) until 99% of the population are double mutants (yielding *σ_DM_* < 1). The dashed line represents the theoretical boundary of the σ*_DM_* value, attained when half of the demes are inhabited solely with double mutants, and the other half is empty of double mutants. The qualitative dynamics are very similar for the different selection coefficients (compare A, B and C) though the scale is different. Other parameters: *n* = 1,000, *k* = 10, μ = 10^−5^, *r* = 0.01, *b* = 1.2.

## References

Andersson, D. I. and D. Hughes. 2010. Antibiotic resistance and its cost: is it possible to reverse resistance? Nature Reviews Microbiology 8:260–271.

Axelrod, R. and W. D. Hamilton. 1981. The evolution of cooperation. Science 211:1390–1396.

Bennett, J. H. 1983. Natural selection, heredity, and eugenics: Including selected correspondence of RA Fisher with Leonard Darwin and others.

Benton, M. J. 2009. The Red Queen and the Court Jester: species diversity and the role of biotic and abiotic factors through time. Science 323:728–732.

Bitbol, A.-F. and D. J. Schwab. 2014. Quantifying the role of population subdivision in evolution on rugged fitness landscapes. PLoS computational biology 10:e1003778.

Caudill, L. and J. R. Wares. 2016. The role of mathematical modeling in designing and evaluating antimicrobial stewardship programs. Current Treatment Options in Infectious Diseases 8:124–138.

Clarke, B. 1979. The evolution of genetic diversity. Proceedings of the Royal Society of London B: Biological Sciences 205.453–474:

Coyne, J. A., N. H. Barton, and M. Turelli. 1997. Perspective: a critique of Sewall Wright’s shifting balance theory of evolution. Evolution:643–671.

Coyne, J. A., N. H. Barton, and M. Turelli. 2000. Is Wright’s shifting balance process important in evolution? Evolution 54:306–317.

de Visser, J. A. G. and J. Krug. 2014. Empirical fitness landscapes and the predictability of evolution. Nature Reviews Genetics 15:480–490.

Eshel, I. 1981. On the survival probability of a slightly advantageous mutant gene with a general distribution of progeny size—a branching process model. Journal of mathematical biology 12:355–362.

Eshel, I. and M. W. Feldman. 1970. On the evolutionary effect of recombination. Theoretical population biology 1:88–100.

Gavrilets, S. 2004. Fitness landscapes and the origin of species (MPB-41). Princeton University Press Princeton, NJ.

Gueijman, A., A. Ayali, Y. Ram, and L. Hadany. 2013. Dispersing away from bad genotypes: the evolution of Fitness-Associated Dispersal (FAD) in homogeneous environments. BMC evolutionary biology 13:125.

Hadany, L. 2003. Adaptive peak shifts in a heterogenous environment. Theoretical population biology 63:41–51.

Hadany, L. and T. Beker. 2003. Fitness-associated recombination on rugged adaptive landscapes. Journal of evolutionary biology 16:862–870.

Hamilton, W. 1964. The genetical evolution of social behaviour. I.

Kagel, J. H. and A. E. Roth. 1995. The handbook of experimental economics. Princeton university press Princeton, NJ.

Kauffman, S. A. and E. D. Weinberger. 1989. The NK model of rugged fitness landscapes and its application to maturation of the immune response. Journal of theoretical biology 141:211–245.

Kingman, J. 1978. A simple model for the balance between selection and mutation. Journal of Applied Probability:1–12.

Komarova, N. L. 2014. Spatial interactions and cooperation can change the speed of evolution of complex phenotypes. Proceedings of the National Academy of Sciences 111:10789–10795.

Kreft, J.-U. 2004. Biofilms promote altruism. Microbiology 150:2751–2760.

Michalakis, Y. and M. Slatkin. 1996. Interaction of selection and recombination in the fixation of negative-epistatic genes. Genetical research 67:257–269.

Nadell, C. D., J. B. Xavier, S. A. Levin, and K. R. Foster. 2008. The evolution of quorum sensing in bacterial biofilms. PLoS biology 6:e14.

Neidhart, J., I. G. Szendro, and J. Krug. 2014. Adaptation in tunably rugged fitness landscapes: the rough Mount Fuji model. Genetics 198:699–721.

Nowak, M. A. 2006. Five rules for the evolution of cooperation. science 314:1560–1563.

Obolski, U., E. Dellus-Gur, G. Y. Stein, and L. Hadany. 2016. Antibiotic cross-resistance in the lab and resistance co-occurrence in the clinic: Discrepancies and implications in E. coli. Infection, Genetics and Evolution 40:155–161.

Obolski, U. and L. Hadany. 2012. Implications of stress-induced genetic variation for minimizing multidrug resistance in bacteria. BMC medicine 10:89.

Obolski, U., G. Y. Stein, and L. Hadany. 2015. Antibiotic Restriction Might Facilitate the Emergence of Multi-drug Resistance. PLoS Comput Biol 11:e1004340.

Ochman, H., J. G. Lawrence, and E. A. Groisman. 2000. Lateral gene transfer and the nature of bacterial innovation. Nature 405:299–304.

Perron, G. G., R. F. Inglis, P. S. Pennings, and S. Cobey. 2015. Fighting microbial drug resistance: a primer on the role of evolutionary biology in public health. Evolutionary applications 8:211–222.

Ram, Y. and L. Hadany. 2014. Stress-induced mutagenesis and complex adaptation. Proceedings of the Royal Society B: Biological Sciences 281:2014–1025.

Salverda, M. L., E. Dellus, F. A. Gorter, A. J. Debets, J. Van Der Oost, R. F. Hoekstra, D. S. Tawfik, and J. A. G. de Visser. 2011. Initial mutations direct alternative pathways of protein evolution. PLoS Genet 7:e1001321.

Thomas, C. M. and K. M. Nielsen. 2005. Mechanisms of, and barriers to, horizontal gene transfer between bacteria. Nature reviews microbiology 3:711–721.

Trivers, R. L. 1971. The evolution of reciprocal altruism. Quarterly review of biology:35–57.

Weinreich, D. M., L. Chao, and P. Phillips. 2005. Rapid evolutionary escape by large populations from local fitness peaks is likely in nature. Evolution 59:1175–1182.

Weissman, D. B., M. M. Desai, D. S. Fisher, and M. W. Feldman. 2009. The rate at which asexual populations cross fitness valleys. Theoretical population biology 75:286–300.

Weissman, D. B., M. W. Feldman, and D. S. Fisher. 2010. The rate of fitness-valley crossing in sexual populations. Genetics 186:1389–1410.

West, S. A,.S. P. Diggle, A. Buckling, A. Gardner, and A. S. Griffin. 2007a. The social lives of microbes. Annual Review of Ecology, Evolution, and Systematics:53–77.

West, S. A., A. S. Griffin, and A. Gardner. 2007b. Evolutionary explanations for cooperation. Current Biology 17:R661–R672.

Whitlock, M. C. 1995. Variance-induced peak shifts. Evolution:252–259.

Whitlock, M. C. 1997. Founder effects and peak shifts without genetic drift: adaptive peak shifts occur easily when environments fluctuate slightly. Evolution:1.044–1048

Williams, S. M. and S. Sarkar. 1994. Assortative mating and the adaptive landscape. Evolution:868–875.

Wright, S. 1932. The roles of mutation, inbreeding, crossbreeding and selection in evolution. Pp. 356–366. Proceedings of the sixth international congress on genetics.

## <u>References</u>

Harris, T. E. 1948. Branching processes. The Annals of Mathematical Statistics:474–494.

